# Hyperactive Rac converts sublethal nibbling to lethal phagocytosis *in vivo*

**DOI:** 10.1101/2025.11.28.691259

**Authors:** Abhinava K Mishra, Lauren Penfield, Morgan Smith, Denise Montell

**Affiliations:** Molecular, Cellular, and Developmental Biology Department University of California, Santa Barbara, CA 93106

**Keywords:** Phagocytosis, Trogocytosis, *Drosophila*, CAR-M, Ovary

## Abstract

The small GTPase Rac is an essential regulator of cell shape, migration, macropinocytosis and phagocytosis. We recently reported that expression of constitutively active Rac^G12V^ is sufficient to cause a few migratory cells called border cells to cannibalize nurse cells in the *Drosophila* ovary. Building on that insight, we engineered mammalian Rac-enhanced chimeric-antigen-receptor macrophages (RaceCAR-Ms) to avidly engulf and kill cancer cells. Here we investigate the cellular and molecular mechanisms by which border cells efficiently kill the much larger nurse cells. Surprisingly, wild type border cells normally nibble on nurse cells as they migrate between them, and Rac^G12V^ causes border cells to take larger, lethal bites. These larger bites trigger rapid germline shrinkage, nuclear damage, and caspase activation, which spreads through the nurse cell syncytium. Then, many somatic follicle cells join in to engulf the dying germline. Rac and the engulfment receptor Draper are critical for both sublethal nibbling and lethal phagocytosis. Using clonal analysis, we show small groups of follicle cells expressing Rac^G12V^ induced caspase activation in neighboring follicle cells while larger Rac^G12V^ clones were required to cause germline killing. Increasing Draper expression or JNK activity in border cells also caused germline death, in a Rac-independent manner, suggesting that border cells can be activated to kill through multiple mechanisms. The series of events elucidated here reveals how hyperactivated Rac expressed in a few cells can trigger destruction of a much larger mass of cells.

**Significance Statement:** Rac is a key protein in the cellular eating process called phagocytosis. Rac hyperactivity enhances the consumption of tumor cells by chimeric antigen receptor-macrophages (CAR-M), a promising type of cellular immunotherapy. Elucidating the mechanisms by which hyperactive Rac enhances cell killing may lead to improvements in CAR-M. Key insights into the *in vivo* effects of Rac have come from studying *Drosophila* oogenesis. Here we report molecular and cellular mechanisms by which hyperactivated Rac stimulates migratory cells to engulf and kill much larger cells in the fly ovary, ultimately resulting in destruction of the entire tissue. These insights have implications for how hyperactivating Rac might improve antibody and CAR-M therapies for cancer and other diseases.

## Introduction

The ability of one cell to eat another arose early in the evolution of eukaryotic cells and is a prevalent behavior in both unicellular and multicellular organisms (1–3). Many protists survive by engulfing smaller cells (4). In animals, macrophages and neutrophils are innate immune cells that specialize in eating and killing pathogens as well as stressed or diseased cells. Macrophage-mediated cell removal is also essential for tissue homeostasis; for example, macrophages remove millions of senescent red blood cells every second. Macrophages also consume dying or dead cells to promote normal development and wound healing and to prevent inflammation. Macrophages eat neutrophils and T-cells to resolve inflammatory responses (5). They remove autoreactive B and T cells to avert autoimmune disease, and macrophage engulfment of incipient tumor cells contributes to cancer prevention. However, in some cases, cancer cells have co-opted this behavior. Tumor cells can feed on T cells, simultaneously suppressing anti-tumor immune attack and scavenging nutrients (6–9).

Chimeric-antigen-receptor (CAR) macrophage therapy harnesses the ability of macrophages to engulf and kill harmful cells (10, 11). Peripheral blood monocytes are collected from cancer patients and engineered to express a CAR that recognizes cancer cells and triggers engulfment. Such CAR-macrophage or CAR-monocyte (CAR-M) immunotherapies are in clinical trials and are safe. However a key challenge is increasing the phagocytic capacity of CAR-Ms (12). In particular, converting sublethal nibbling, or trogocytosis, (13) to lethal phagocytosis is a major goal for the field (14). Therefore, enhancing engulfment of living target cells in native, 3D environments is essential. One way to learn more about how cells engulf and kill other living cells *in vivo* is to study such behaviors in relatively simple experimental organisms like the fruit fly *Drosophila* (15, 16).

Normal *Drosophila* oogenesis occurs in structures called egg chambers, which grow through 14 developmental stages as they mature. Egg chambers contain somatic follicle cells surrounding 16 germ cells, including 15 support cells called nurse cells and one egg cell (oocyte) (Fig. 1A). The germ cells are interconnected via cytoplasmic bridges called ring canals, through which the nurse cells donate cytoplasm to the oocyte. Afterwards, nurse cells are no longer needed, and so-called stretch follicle cells remove them (17) (Fig. 1B). Expression of engulfment genes in stretch follicle cells is required for nurse-cell death and consumption (18). The engulfment machinery includes transmembrane proteins such as integrins and the scavenger receptor Draper (the fly homolog of human MEGF10/11/12), as well as intracellular signaling proteins like the kinases Src, Shark (Drosophila Syk), and Jun N-terminal kinase (JNK).

**Fig. 1.**
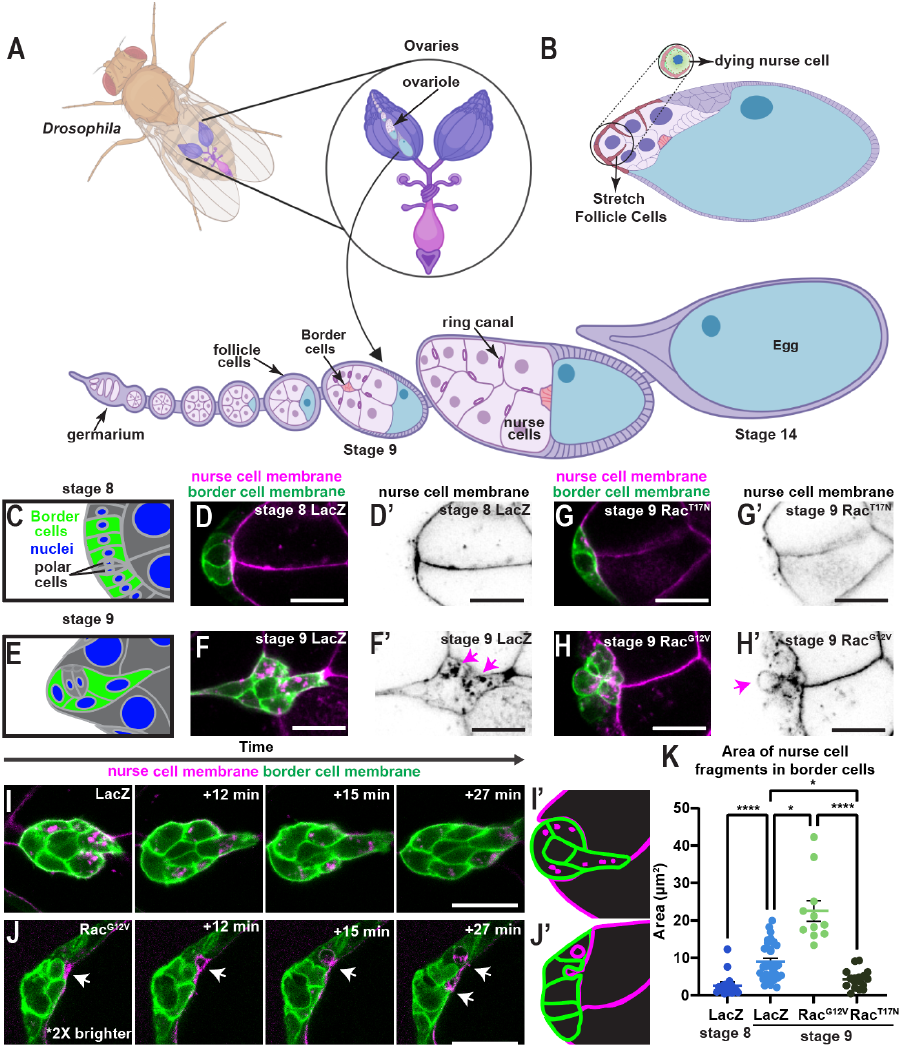
Wild type border cells nibble on nurse cells and Rac^G12V^ border cells take larger bites. (A) Schematic of *Drosophila* ovaries and egg chamber stages. Border cells migrate in stage 9. (B) In stage 12, stretch cells begin to engulf nurse cells. (C) Schematic of border cells at stage 8. (D) Confocal image of control stage 8 border cell cluster expressing *slbo*4x-PH-EGFP (green). Nurse cell membranes (magenta) are labeled with UMAT-Lyn-TdTomato. D’ shows inverted fluorescence of maternal driven UMAT-Lyn-TdTomato (black). (E) Schematic of delaminating border cell cluster at stage 9. (F) Confocal image of control stage 9 border cell cluster expressing *slbo*4X-PHE-GFP and nurse cell membranes are labeled with UMAT-Lyn-TdTomato F’. inverted fluorescence image of UMAT-Lyn-TdTomato (G-H) Images of stage 9 border cells labeled with *slbo*4x-PH-EGFP (green). Nurse cell membranes are labeled with UMAT-LynTd-Tomato (magenta). *slboGal4* drives *UAS-Rac*^*T17N*^ (G-G’) or *UAS-Rac*^*G12V*^ (H-H’). (G’-H’) Inverted fluorescence of UMAT-Lyn-TdTomato (black). Magenta arrows point to engulfed UMAT-LynTdTomato. (I-J) Images from live confocal imaging of border cells expressing *slboGal4* with *UAS-LacZ slbo*4x-PH-EGFP, UMAT-Lyn-TdTomato (G), or *UAS-Rac*^*G12V*^ *slbo*4x-PH-EGFP, UMAT-Lyn-TdTomato. Rac^G12V^ egg chambers had dimmer fluorescence, the images have been adjusted differently to match signal brightness. White arrows indicate UMAT-Lyn-TdTomato inside of border cells. See also Movies S1 and S2. (K) Maximum area of nurse cell fragment within border cells for the indicated genotypes and stages. All genotypes included *slboGal4; UMAT-Lyn-TdTomato*. Mean +/-standard error of the mean (SEM). Each dot represents a single egg chamber (n>10) from 3 or more experimental replicates. Statistical analysis is from a Kruskal Wallis test. ^*^ p<0.05, ^****^ p<0.0001. Scale bars: 20 μm.

In addition to this late-stage killing and removal of nurse cells, protein-deprived flies redirect nutrients from egg production to survival. This nutrient checkpoint occurs earlier, at stage 7/8 of egg development (19, 20). During nutrient stress, nurse cells undergo apoptosis, then all the surrounding follicle cells engulf the dying germline. Developmental nurse cell removal and starvation-induced egg chamber destruction employ some overlapping and some distinct mechanisms (17, 19).

We recently discovered that expressing an activated form of the small GTPase Rac, which is generally required for phagocytosis (21), in a few migratory follicle cells called border cells is sufficient to destroy the entire egg chamber (15). Building on this insight, we showed that hyperactivate Rac is also sufficient to cause mouse and human macrophages to engulf and kill chronically activated T-cells, likely contributing to T-cell loss in patients with dominant, activating Rac2 mutations. Moreover, this hyper engulfment behavior can be harnessed to enhance CAR-M-mediated cancer cell killing (15).

Here we report the cellular and molecular mechanisms by which Rac stimulates cell engulfment and killing of Drosophila egg chambers. We discovered that border cells normally nibble on germline nurse cells as they migrate between them. Germline nibbling requires the engulfment receptor, Draper, and Rac. RacG12V-expressing border cells take larger bites, triggering nurse cell shrinkage followed by loss of nuclear integrity, caspase activation and apoptotic cell death, which spreads rapidly through the nurse cell syncytium. Other follicle cells join in to consume the dying germline. Together these data show that Rac hyperactivity converts normal, sublethal nibbling of nurse cells to lethal trogocytosis, a major goal in CAR-M cancer immunotherapy.

## Results

### Wild type border cells nibble on nurse cells and Rac^G12V^ border cells take larger bites

The *Drosophila* ovary is composed of strands of developing egg chambers of increasing size and maturity (Fig. 1A). Germline and somatic stem cells reside at the anterior tip in a structure known as the germarium. Their progeny assemble into ovoid egg chambers composed of a syncytium of 16 interconnected germline cells surrounded by epithelial follicle cells. The most posterior of the germ cells differentiates into the oocyte, while the rest become polyploid nurse cells. As development progresses, the germline grows and the oocyte occupies an increasing percentage of the egg chamber. At developmental stage 8, anterior follicle cells termed polar cells induce 4-8 neighboring cells to become border cells. At stage 9, the border follicle cells delaminate from the epithelium and migrate collectively (Fig. 1A), eventually reaching the anterior border of the oocyte by stage 10 (22). At stage 11, the nurse cells transfer the majority of their cytoplasm to the oocyte, which rapidly enlarges. At stage 12, anterior follicle cells termed stretch cells surround, isolate, kill, and engulf each nurse cell (Fig. 1B) (17).

The observation that Rac^G12V^ expression in a few border cells is sufficient to kill the entire egg chamber (15) raises the question as to how a handful of small cells can kill much larger cells. Draper contributes to Rac^G12V^-mediated egg chamber death because mutating Draper rescues about 50% of the egg chamber death (15), implying that Rac^G12V^-mediated killing requires engulfment. To visualize this process directly, we acquired high-magnification images of egg chambers in which nurse cell membranes were labeled with a maternally driven membrane targeting sequence of Lyn fused to Tdtomato (UMAT-LynTdTomato) (23). Border cell membranes were labeled with a plextrin homology (PH) domain from phospholipase C δ (PLCδ) fused to enhanced green fluorescent protein (*slbo*4x-PH-EGFP) expressed in border cells using an enhancer derived from the *slow border cells* (*slbo*) gene (24). We expressed *UAS-Rac*^*G12V*^ or a control (*UAS-LacZ*) in border cells with a border cell Gal4 driver, *slboGal4* (24). While we expected nurse cell membranes to be internalized into Rac^G12V^-expressing border cells, we were surprised to see red fluorescence accumulating within control (*slboGal4;UAS-LacZ*) border cells (Fig. 1C-F). At stage 8, a few small puncta were visible within border cells (Fig. 1D and D’). However at stage 9, as border cells began migrating, they accumulated larger nurse cell membrane fragments (Fig. 1F and F’). Thus, border cells normally nibble as they move between nurse cells. This nibbling was dependent on Rac activity because expressing dominant-negative Rac (Rac^T17N^) prevented the increase at stage 9 (Fig. 1G and G’). The reduced nibbling by Rac^T17N^-expressing border cells was not a secondary consequence of their inability to migrate because Rac^G12V^-expressing border cells also fail to move but accumulated nurse cell membrane puncta (Fig. 1H and H’). Control border cells contained small puncta throughout the migration (Fig. 1I and I’ and Movie S1) whereas Rac^G12V^ border cells engulfed larger fragments of nurse cell membranes over time (Fig. 1 J and J’ and Movie S2). Thus, expression of hyperactive Rac^G12V^ in border cells increases the sizes of engulfed fragments (Fig. 1K).

### The engulfment receptor Draper is required for border cell nibbling and migration

Since control border cells nibble on nurse cells, we investigated the expression and function of Draper in border cells. Draper expression increased in border cells compared to other follicle cells, specifically at stage 9 when nibbling increased (Fig. 2A-D). Then, we evaluated border cell nibbling in *drpr* mutants, which are homozygous viable. In control egg chambers, 100% of border cell clusters complete migration by stage 10 (Fig. 2E) (25). In the *draper* mutant, 60% of clusters failed to complete migration by stage 10 (Fig. 2F and G), consistent with a previous report (26). Thus, Draper is required for efficient border cell migration.

**Fig. 2.**
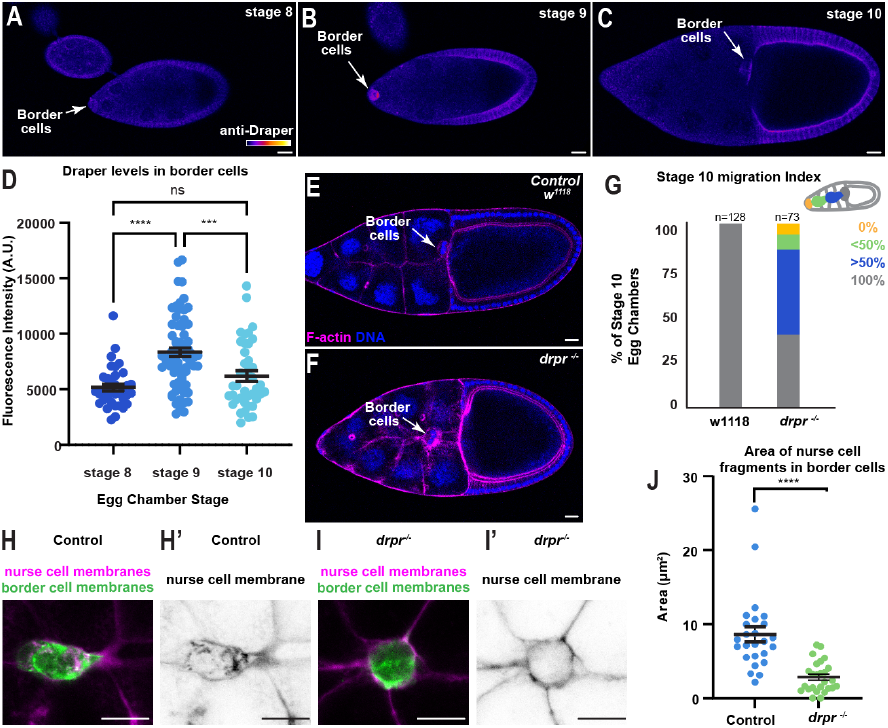
Draper promotes nurse cell nibbling and migration in border cells. (A-C). Anti-Draper antibody staining in wild type stage 8-10 egg chambers. Draper expression shown in Fire LUT scale (Image J) shown. (D) Quantification of Draper levels at the indicated egg chamber stages. Bars show mean +/-SEM. Each dot (n>40) represents a border cell cluster from 3 independent dissections and stainings of *w*^*11118*^, *Sco/CyO; MRKS/TM6B* flies. ^***^ p<0.001, ^****^ p<0.0001. (E-F) Images of control (*w*^*11118*^, *Sco/CyO; MRKS/TM6B*) and Draper mutant (*drpr-/-*) egg chambers stained with phalloidin (magenta) and Hoechst (blue). All border cell clusters reach the oocyte by stage 10 in control but not in *drpr* mutants. (G) Quantification of border cell migration completion in control vs. *drpr* mutants, n=number of egg chambers from 8 (*drpr-/-*) or 9 (control) experimental replicates. (H-I) Confocal images of border cell membranes marked with *slbo*4X-PH-EGFP (green) and nurse cell membranes marked with UMAT-LynTdTom (magenta in H and I, black in H’ and I’) in control (H) and *drpr* mutant (I) egg chambers. (J) Maximum area of a nurse cell membrane fragment within a border cell cluster for the indicated genotypes =. Mean +/-standard error of the mean (SEM). Each dot represents an egg chamber n>25 egg chambers from 3 or more experimental replicates. ^****^p<0.0001. Scale bars: 20 μm.

We next assessed if Draper is required for border cell nibbling. Border cells in *draper* mutants engulfed significantly less nurse cell membrane material compared to control border cells (Fig. 2 H-J). We conclude that border cells normally increase Draper-and Rac-dependent nibbling in stage 9, as they migrate, without killing the germline.

Together, these data suggest that the Rac^G12V^-induced effect on germ cells is an exaggeration of normal nibbling rather than a completely unnatural effect and indicate that increased Rac activity is sufficient to convert sublethal nibbling into lethal engulfment. We then explored the sequence of events that result in germline killing.

### Border cells expressing Rac^G12V^ induce germline chromatin condensation throughout the syncytium

To test if Rac^G12V^ nibbling preceded egg chamber death, we imaged egg chambers with border cells expressing a membrane marker and nurse cells expressing an Ubiquitin driven Histone fused to RFP (Ubi-His-RFP) in nuclei and UMAT-Lyn-TdTomato at plasma membranes (Fig. 3A and Movie S3). In control egg chambers, border cells migrate between nurse cells, whereas Rac^G12V^-expressing border cells do not migrate (15) (Fig. 3A and B). We found that control nurse cell nuclei contained relatively uniformly distributed His-RFP (Fig. 3A and Movie S3), but when border cells expressed Rac^G12V^, nurse cell nuclei underwent chromatin condensation (Fig. 3B’ and B’’ and Movie S4). Higher magnification imaging showed that border cells normally extend protrusions between nurse cells but do not touch or perturb the nuclei (Fig. 3C, Movie S5). However, Rac^G12V^ border cells extend protrusions that pinch both nurse cell membranes and nurse cell nuclei prior to chromatin condensation (Fig. 3D and E, and Movies S6 and S7). In some examples, chromatin changes occurred in all nurse cells within the same 3-minute imaging interval (Fig. 3B); however, we also observed examples where chromatin condensation clearly initiated in one cell that was interacting with a border cell and then spread rapidly to the other cells (Fig. 3 E and F). These data indicate that Rac^G12V^ border cells trigger germline death, which rapidly spreads through the syncytium.

**Fig. 3.**
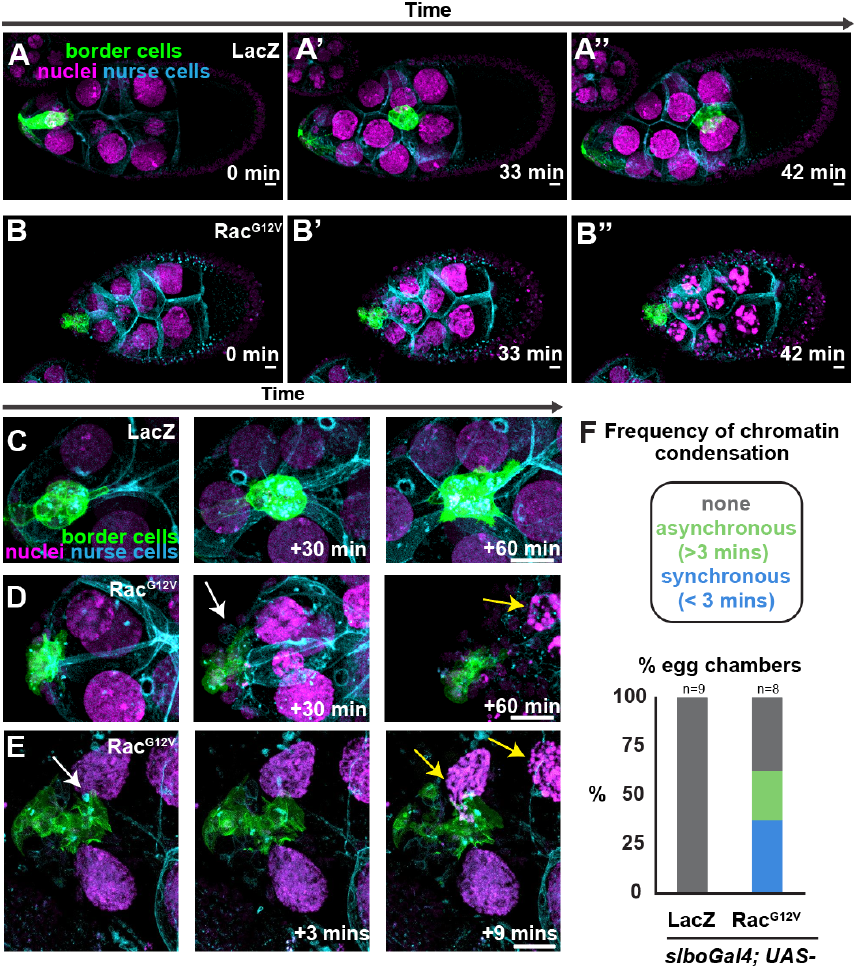
Rac^G12V^ border cells induce a rapid death of nurse cells that spreads through the syncytium. (A-B) Images from timelapse movies of control (*slboGal4; UAS-LacZ*) border cells migrating or Rac^G12V^ border cells (*slboGal4;UAS-Rac*^*G12V*^). Border cells are labeled with *slbo*-4xPHEGFP, nuclei are labeled with UbiHisRFP, and membranes are labeled with UMAT-Lyn-TdTomato. Scale bars: 10 μm. Images were acquired by Lambda imaging and channels were distinguished by spectral unmixing. See also Movies S3 and S4. (C-E) Images from a timelapse series from higher magnification imaging of control and Rac^G12V^-expressing border cells with the same fluorescent markers. White arrows note membrane pinching, yellow arrows indicate chromatin condensation. See also movies S5-7. Scale bars: 20 μm. Images were acquired by Lambda imaging and channels were distinguished by spectral unmixing. (F) Plot showing the frequency of germline death as indicated by chromatin condensation in control and Rac^G12V^ imaging sessions. n>8 egg chambers per condition. Germline death was scored as synchronous if all nuclei condensed with similar dynamics within a 3 minute image acquisition interval, and asynchronous if they condensed at different rates or times.

### Rac^G12V^-mediated engulfment triggers nurse cell shrinking prior to caspase-dependent chromatin condensation

To investigate whether nurse cells undergo apoptotic cell death, we evaluated caspase activation by staining for the cleaved and activated form of Dcp-1 (cDcp-1), an executioner caspase. The majority of *slboGal4;UAS-Rac*^*G12V*^ egg chambers with condensed chromatin were positive for cDcp-1 (Fig. 4A-C). Active caspases cleave nuclear lamins and other components of the nuclear envelope, resulting in loss of the nuclear permeability barrier (27, 28). Live imaging of control stage 9 egg chambers with an Ubiquitin promoter driving NLS-GFP showed that control egg chambers retained the nuclear permeability barrier over time (Fig. 4D and D’ and Movie S8) while in egg chambers with Rac^G12V^ border cells revealed that the nuclear permeability barrier was lost in a subset of germline cells and rapidly spread to neighboring cells (Fig. 4E and E’’ and Movie S9). In contrast, we did not observe signs of lytic cell death such as nurse cell membrane rupture. 10 kDa dextrans accumulate in intercellular junctures between nurse cells in control (Fig. S1A) as previously reported (29). In egg chambers with Rac^G12V^ border cells, 10 kDa dextrans also accumulate between nurse cells, but are excluded from the nurse cell syncytium despite chromatin condensation (Fig. S1B). These data indicate that the large Rac^G12V^-mediated bites trigger apoptosis rather than necrosis within the nurse cell syncytium.

**Fig. 4.**
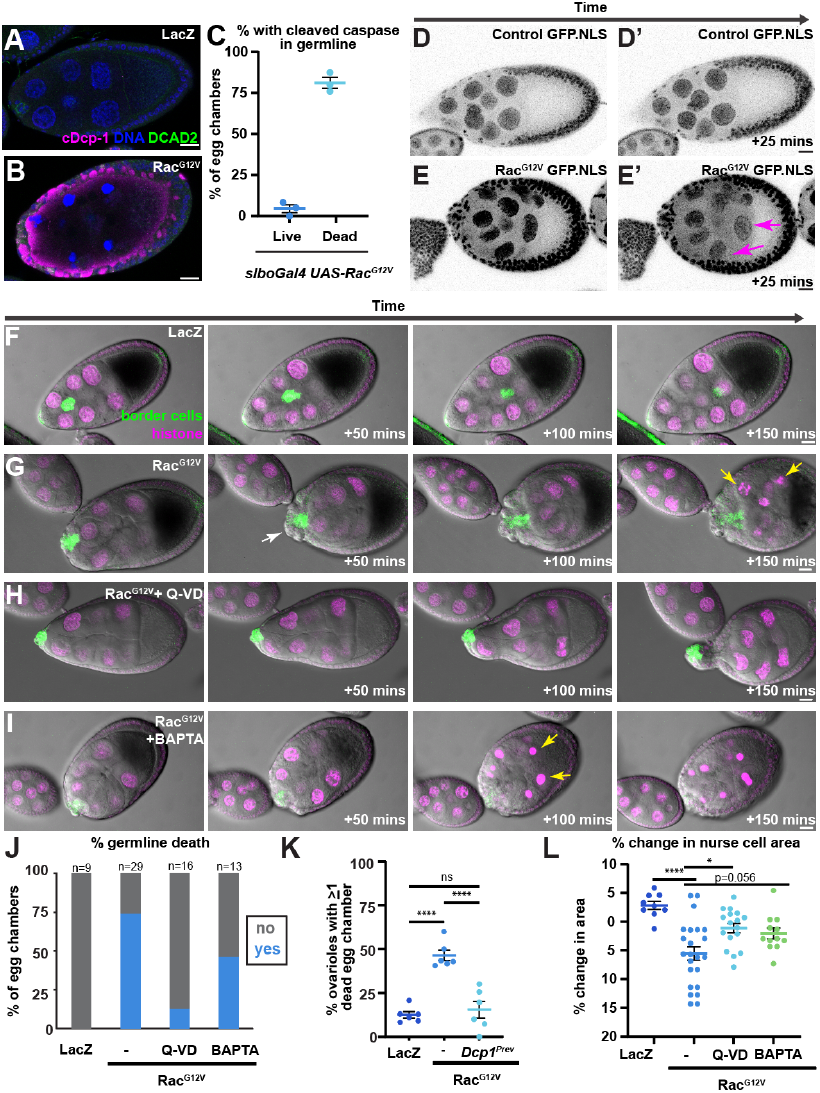
Rac^G12V^-mediated engulfment triggers nurse cell shrinkage prior to caspase-dependent chromatin condensation. (A-B) Representative confocal images of egg chambers with the indicated conditions and markers. Anti-cleaved caspase (cDcp-1) is shown in magenta. (C) Plot showing the percent of egg chambers that stained positively for cDcp-1 in the germline for the indicated conditions analyzed from 15 or more egg chambers from 3 independent experiments. (D-E’) Images from timelapse series of egg chambers expressing Ubiquitously expressed GFP.S65T with a Nuclear Localization Signal (Ubi-NLS, green) was used to assess the nuclear permeability barrier in control egg chambers (D-D’) and egg chambers with *slboGal4; UAS-Rac*^*G12V*^ (E-E’) See also Movies S8 and S9. F-I Images from time lapse series of egg chambers for the indicated conditions. Genotypes/treatments are *slboGal4;* UbiHisRFP, *slbo*4XPH-EGFP with *UAS-LacZ* (F), *UAS-Rac*^*G12V*^ (G) *UAS-Rac*^*G12V*^ +100 micromolar Q-VD-OPh (H) or *UAS-Rac*^*G12V*^ + 50 micromolar BAPTA-AM (I). White arrows indicates egg chamber with germline shrinking; yellow arrows denote chromatin condensation. Scale bars 20 μm See also Movies S10-13. (J) Percentage of egg chambers with condensed germline chromatin within the 4.5 hour imaging session. n=number of egg chambers, from N=2 (LacZ) or 4 experimental replicates (*UAS-Rac*^*G12V*^ +/-drug treatments). (K) Percentage of egg chambers with condensed germline chromatin in ovarioles from fixed samples for the indicated genotypes. Mean +/-SEM (bars). Each dot represents one experiment where 14 or more egg chambers were analyzed. Data are combined from at least 3 separate crosses. One-way ANOVA followed by a post hoc Tukey test. (L) Percent change in nurse cell area over 25 minutes (see methods). Mean +/-SEM for the indicated genotypes and treatments. Each dot represents the average of the change between time points in a movie, prior to chromatin condensation, for a single egg chamber, n> 9 egg chambers from 2 (LacZ) or 4 (Rac^G12V^) experimental replicates. One way ANOVA followed by a post hoc Tukey test. ^*^ p<0.05, ^***^p<0.001.

Cell shrinkage is a defining characteristic of apoptotic cells and distinguishes it from necrotic cell death. Apoptotic volume decrease can occur before caspase activation (30, 31). Egg chambers normally enlarge during stage 9 (Fig. 4F and Movie S10), so it was striking that *slboGal4;UAS-Rac*^*G12V*^ egg chambers typically shrank after nurse cell membrane internalization and prior to chromatin condensation (Fig. 4G and Movie S11). Consistent with the idea that nurse cells undergo apoptotic volume decrease prior to caspase activation, we also noticed an anteriorward flow of stretch follicle cells during shrinkage (Fig 4G and Movie S11). Interestingly, stretch follicle cells start out as cuboidal and have been proposed to flatten out during stage 9 as a consequence of increasing germline volume (32), somewhat like a balloon stretching as it is inflated. Volume decrease then leads to unstretching, like a deflating balloon. While we considered the possibility that a Ca^2+^ influx that typically occurs upon plasma membrane damage might induce acto-myosin contractility in the nurse cells, we were unable to detect changes in calcium influx (Fig. S1C).

To further test possible contributions of caspase and/or calcium influx, we treated egg chambers with Rac^G12V^-expressing border cells with a pan-caspase inhibitor Q-VD-OPh (herein Q-VD), (Fig. 4H) or BAPTA-AM, a cell permeable calcium chelator (Fig. 4I). Within a 4-hour imaging time frame, Q-VD significantly reduced the frequency of germline chromatin condensation despite some germline shrinkage and nuclear deformations (Fig. 4I and J and Movie S12). As an additional approach, we expressed Rac^G12V^ in border cells in homozygous loss of function caspase mutant *Dcp-1*^*Prev*^. Although we observed a significant reduction in egg chamber death (Fig. 4K), the egg chambers lacked detectable follicle cells including border cells, making the rescue difficult to interpret (Fig. S2). The absence of follicle cells in *Dcp-1*^*Prev*^ egg chambers has been previously noted (33). Interestingly, *slboGal4;UAS-Rac*^*G12V*^ egg chambers underwent some morphological changes even in the presence of Q-VD, so we assessed how germline shrinking was affected. Q-VD treatment significantly reduced the germline shrinking in *slboGal4;UAS-Rac*^*G12V*^ egg chambers, yet it did not fully restore growth (Fig. 4L). We conclude that slboGal4;UAS-Rac^G12V^ induced nibbling and germline shrinkage, followed by caspase-dependent chromatin condensation and further shrinkage.

Upon treatment with BAPTA-AM, Rac^G12V^-expressing border cells killed 46% egg chambers examined by live imaging (Fig. 4I and Movie S13), compared to 79% of egg chambers killed without BAPTA-AM (Fig. 4J). Further BAPTA-AM reduced the extent of germline shrinkage (Fig. 4I and L). These data indicate intracellular calcium contributes to Rac-mediated germline death. We propose that as border cells take bites out of the anterior nurse cells, it triggers apoptosis characterized by early volume decrease followed by caspase activation and chromatin condensation.

### Rac^G12V^-induced death is distinct from both starvation-induced and developmental germline death

There are two situations in which follicle cells engulf the germline. There is a checkpoint at stage 7/8, and if females are protein-restricted, egg chambers do not develop further. Rather, the germline activates caspase, undergoes apoptosis and is engulfed by follicle cells, to redirect nutrients toward survival rather than reproduction (19). In protein-rich conditions, egg chamber development proceeds, nurse cells transfer their cytoplasm to the oocyte during stage 11, and as they shrink, the neighboring stretch follicle cells kill them, in a process that depends on engulfment machinery (17).

In starvation mediated death, follicle cells begin to engulf the germline once it undergoes apoptosis, which we could visualize by uptake of nurse cell membranes (UMAT-LynTd-Tomato) (Fig. 5A and A’). In developmental death, the stretch cells engulf the germline at stage 12 (Fig. 5B and B’). In control stage 8 egg chambers, many follicle cells contained small puncta of UMAT-LynTdTom (Fig. 5C and C’), which only noticeably increased at stage 9 within the border cells (Fig. 5D and D’). In egg chambers expressing Rac^G12V^ in border cells, puncta in the majority of follicle cells remained small in stage 8 (Fig. E-E’) and some intact stage 9 egg chambers (Fig. 5F-F’). However, in shrunken and rounded stage 9 egg chambers, additional follicle cells increased their uptake of nurse cell membranes (Fig. 5G-H), similar to starved egg chambers (Fig. 5A). We conclude that Rac^G12V^ border cells initiate germline death, which then triggers other follicle cells to join in to remove dying germ cells.

**Fig. 5.**
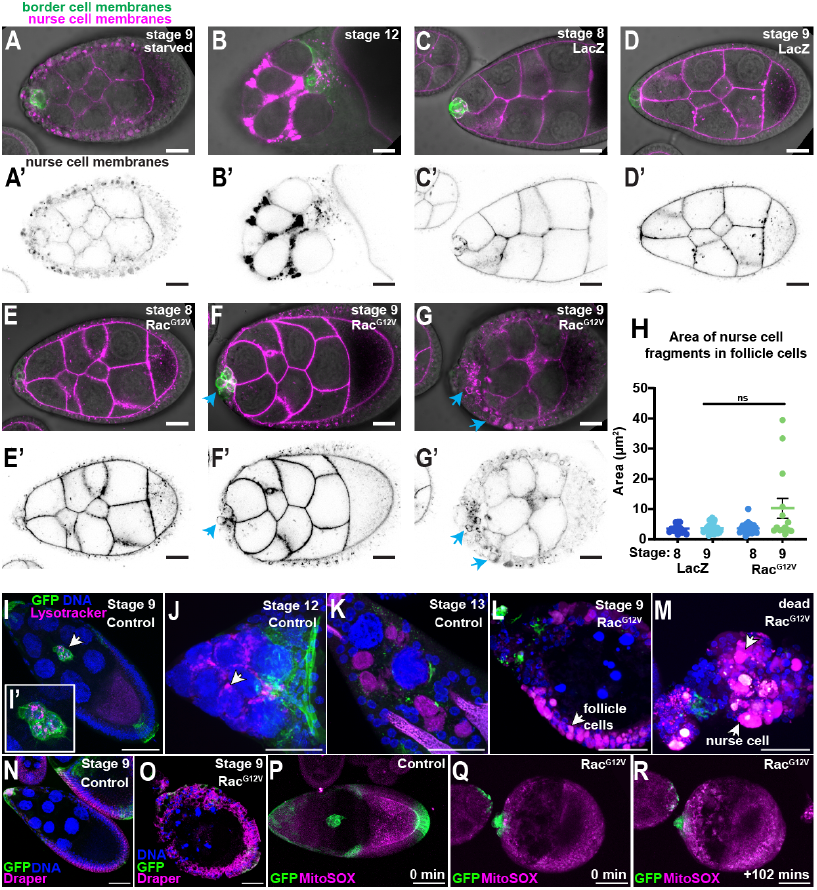
Rac^G12V^ mediated death is distinct from starvation and developmental mediated death. (A-G) Confocal images of egg chambers in Differential interference contrast (DIC) microscopy and fluorescence of border cell membranes marked with *slbo*4XPH-EGFP (green) and nurse cell membranes marked with UMAT-LynTdTomato (magenta) in egg chambers protein starved overnight or in (A) control stage 12 (B) control stage 8 (C) control stage 9 (D) egg chambers in control, yeast supplemented flies. E-G. show *slboGal4; UAS-RacG12V* egg chambers at stage 8 (E), stage 9 (F), and upon death (G). (A’-G’) inverted fluorescence of UMAT-LynTd-Tomato. Scale bars: 20 μm. (H) Mean +/-SEM of the area of nurse cell membrane fragment in follicle cells for the indicated conditions. Each dot represents the area of the largest fragment within follicle cells of an individual egg chamber (n>10) from 3 or more experimental replicates. Significance was tested with a Kruskal-Wallist test, ns=not significant (p>0.05). (I-M) Select confocal images of fixed egg chambers stained with lysotracker (magenta), GFP (green) and DNA (blue) in control stage 9 (I) and magnified image of the border cells (I’), control stage 12 (J), control stage 13 (K), *slboGal4; UAS-Rac*^*G12V*^ stage 9 (L), and later stage 9 in *slboGal4; UAS-Rac*^*G12V*^ (M). (N-O) Confocal images of egg chambers in control stage 9 (N) or egg chambers expressing *slboGal4; UAS-Rac*^*G12V*^ *UAS-LifeActGFP*. (O) stained with mouse-anti Draper (magenta), Hoechst (blue). P-R. Images from timelapse series *slboGal4; UAS-PLCδGFP* egg chambers expressing a control construct *UAS-LacZ* (P) or expressing *UAS-Rac*^*G12V*^ (Q-R). Scale bars: 50 μm (I-R).

During normal development, caspase activation does not initiate nurse cell death. Instead, stretch follicle cells surround and isolate individual nurse cells and secrete lysosomal enzymes, creating acid baths that degrade each nurse cell asynchronously (18). To test if follicle cells also acidify nurse cells during Rac-mediated death we used Lysotracker dye, which fluoresces within acidic compartments. In control stage 9 egg chambers, border cells exhibit larger lysotracker-positive particles than other follicle cells (Fig. 5I, Fig. S3A-C), consistent with their nibbling behavior. At stages 12-13, abundant Lysotracker staining becomes evident first surrounding nurse cells (Fig. 5J), then within nurse cells (Fig. 5K). In *slboGal4;UAS-Rac*^*G12V*^ egg chambers with germline chromatin condensation, most follicle cells had strong lysotracker staining (Fig. 5L), as they engulfed and degraded nurse cell material. Lysotracker accumulated in germline cells only late in the death process (Fig. 5M). We conclude that acidification is a late event in Rac^G12V^-induced germline death.

We performed live imaging to determine changes in lysotracker over time. In controls, we detected lysotracker-positive puncta in border cells (Fig. S3A-B’ and Movie S14) and faint lysotracker signal in other follicle cells (Fig. S3C and C’), which did not change substantially over time (Fig S3A and Movie S14). In egg chambers with Rac^G12V^-expressing border cells, lysotracker accumulated brightly around and in the border cells (Fig. S3D-E’ and Movie S15) and also accumulated in puncta in distant follicle cells (Fig. S3F-F’). Lysotracker positive puncta in follicle cells became larger over time even prior to chromatin fragmentation (Fig S3D, +75 mins, and Movie S15). Nurse cells contained small lysotracker-positive puncta but there was no detectable change in lysotracker upon chromatin condensation (Fig. S3D, +177 min and Movie S15). These data suggest that acidification of the nurse cells does not initiate germline death.

We next assessed Draper levels in control and *slboGal4;UAS-Rac*^*G12V*^ egg chambers. In starvation-mediated death, all follicle cells increase Draper levels after germline nuclei begin to condense (34), while in developmental death, stretch cells specifically increase Draper prior to nurse cell death (26). Draper is normally expressed in all follicle cells (Fig. 5N), as previously reported (34). Draper levels increased in all follicle cells in *slboGal4;UAS-Rac*^*G12V*^ egg chambers with condensed germline chromatin (Fig. 5O), as they engulfed and consumed the dying cells.

Draper overexpression in all follicle cells induces oxidative stress in anterior follicle cells (35), so we assessed the effect of border cell Rac^G12V^ expression on superoxide production using a mitochondrial superoxide sensor, MitoSOX. While there was some staining in control egg chambers (Fig. 5P and Movie S16), MitoSOX increased in follicle cells and in the germline cells over time in *slboGal4;Rac*^*G12V*^ egg chambers, after the egg chambers shrank (Fig. 5Q and R, Movie S17), concomitant with increased Lysotracker and Draper labeling and consumption of the dead germline. We conclude that Rac-mediated death shares features of developmental and starvation-mediated death but is a distinct process as summarized in Table 1.

**Table 1.**
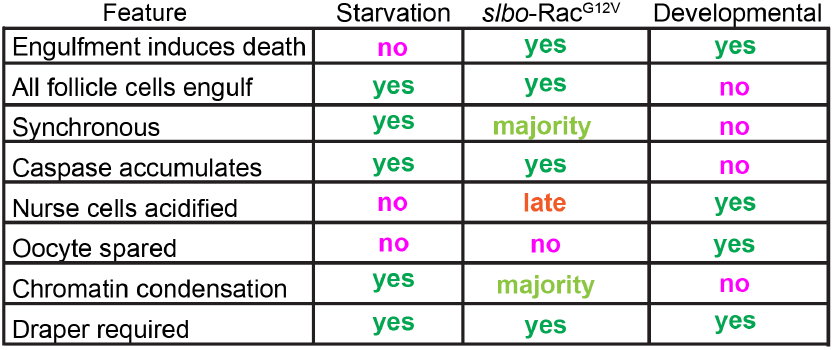
Features of germline death.

| Feature | Starvation | *slbo-RacG12V* | Developmental |
| --- | --- | --- | --- |
| Engulfment induces death | no | yes | yes |
| All follicle cells engulf | yes | yes | no |
| Synchronous | yes | majority | no |
| Caspase accumulates | yes | yes | no |
| Nurse cells acidified | no | late | yes |
| Oocyte spared | no | no | yes |
| Chromatin condensation | yes | majority | no |
| Draper required | yes | yes | yes |

### Border cell Draper-JNK signaling is sufficient to induce germline death

JNK is critical for germline engulfment in response to starvation (34), developmental nurse cell death (26), and Draper-induced germline death (35), so we investigated the role of JNK in Rac^G12V^-induced death. We used *puc*-LacZ, a JNK activity reporter, and an antibody that recognizes the phosphorylated, active form (pJNK) to analyze JNK in Rac^G12V^ expressing cells. Combined with our previous result that Draper is required for about half of the Rac-mediated death (15), Border cells normally exhibit increased *puc-*LacZ staining (Fig. 6A) suggesting elevated JNK activity (36). In *slboGal4;UAS-Rac*^*G12V*^ egg chambers, a few additional follicle cells show increased *puc*-lacZ (Fig. 6B), and more follicle cells gained JNK activity as the germline died (Fig. 6C-D). To measure JNK activity in cells with or without Rac^G12V^ expression, we compared phosphorylated JNK(pJNK) in follicle cell clones with or without Rac^G12V^ expression. Rac^G12V^ expression resulted in a 1.2 fold increase in pJNK levels (Fig. 6E-H), which seems insufficient to account for Rac^G12V^ induced death.

**Fig. 6.**
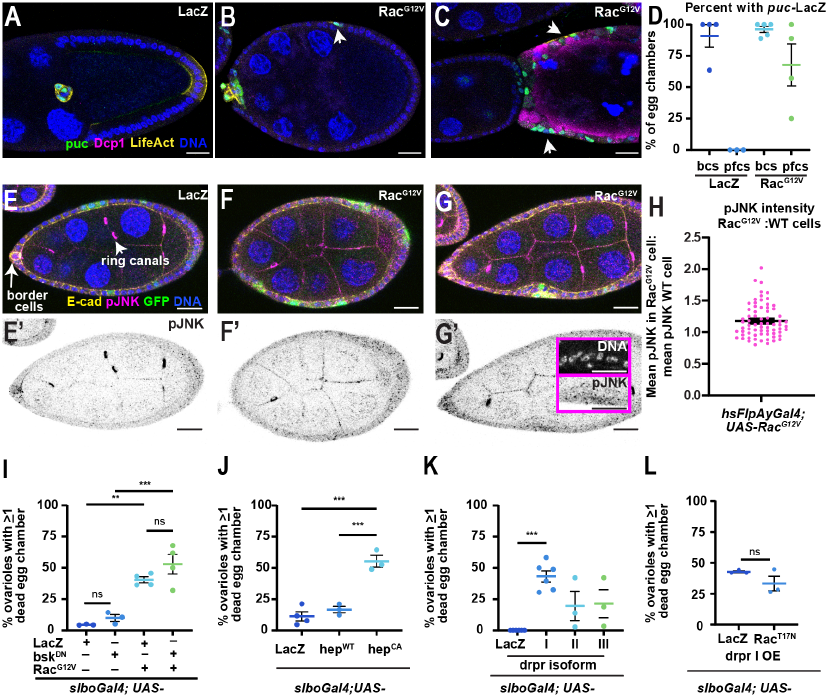
Increasing Draper-JNK signaling in border cells is sufficient to induce germline death. (A-C) Confocal images of control (*slboGal4; UAS-LifeActRFP, puc-*LacZ) or egg chambers with *slboGal4; UAS-LifeActRFP, UAS-Rac*^*G12V*^*puc-*LacZ (B-C). Arrowheads denote *puc*-LacZ positive cells. (D) Plot showing percent of cells with *puc*-LacZ. Bc: border cells pfc: posterior follicle cells. Each dot represents the percent of positive egg chambers (9 or more) from one experimental replicate. (E-G’) Images of FlpOut Gal4 clones where *hsFlpAyGal4* drives *UAS-LacZ* control (E) or *UAS-Rac*^*G12V*^(F-G) along with *UAS-GFP* to mark positive clones. pJNK (magenta) in merged images and (E’-G’) inverted greyscale of pJNK. (H) Plot showing mean +/-SEM fold change in pJNK levels in Rac^G12V^, GFP+ versus wildtype (GFP-) cells, each dot represents one clonal cell that are combined from 3 experimental replicates of 10 or more clones analyzed. One sample Wilcoxon rank test where median is assumed to be 1 give a significant result, ^****^ p<0.0001. (I-L) Plots showing the mean +/-SEM percent of ovarioles strands with dying egg chambers for indicated conditions. UAS-LifeActRFP was also expressed in each line. Each dot represents an experimental replicate with 18 or more egg chambers scored per condition. One way ANOVA with Post-Hoc Tukey (I-K) or a Mann-Whitney test (L) were used to test statistical significance. ^***^p<0.001, ^**^p<0.01, ns=not significant (p>0.05)

To test if JNK activity was required for border cells to induce death, we co-expressed the dominant negative (DN) form of JNK (*UAS-Bsk*^*DN*^) or a control (*UAS-LacZ*) with *UAS-Rac*^*G12V*^ in border cells using *slbo*Gal4 or another border cell driver, *c306Gal4*. Egg chamber death was not significantly reduced in border cells expressing Rac^G12V^ with Bsk^DN^ compared to the LacZ control in either case (Fig. 6I and S4A). We conclude that JNK activity is not required in border cells for Rac^G12V^ to induce germline death.

To test if excess JNK activity was sufficient to induce death when expressed in border cells, we expressed a constitutively active form of the JNK kinase (hep^CA^) with *slboGal4* or *c306Gal4;Gal80*^*ts*^. Expression of hep^CA^ but not wild type JNK kinase (hep^WT^) induced significant germline death (Fig. 6J and Fig. S4A). Similarly, overexpression of Draper isoform I, the only isoform that contains the immunoreceptor tyrosine-based activation motif (ITAM), was sufficient to induce a significant increase in egg chamber death in ovarioles (Fig. 6K, Fig. S4B); however death occurred later in egg chamber development than hep^CA^-or Rac^G12V^-induced death. Upon Drpr I overexpression, the majority of egg chambers survived past stage 8, which was not the case for hep^CA^ or Rac^G12V^ expression driven with *c306Gal4* (Fig. S4C). To test if the Draper-JNK mediated death required RAC activity, we expressed a dominant negative form of Rac^T17N^ along with Draper with *slboGal4*. Rac^T17N^ did not significantly rescue egg chamber death induced by Draper overexpression (Fig. 6L). These data indicate there are parallel pathways in border cells that can induce germline death.

### Rac^G12V^ expression in non-border follicle cells also causes egg chamber death

To compare the effects of *UAS-Rac1*^*G12V*^ in non-border follicle cells to border cells, we used *PG150-Gal4* to drive expression in stretch cells, the cells that normally engulf nurse cells late in development (17) (Fig. 7A). PG150-driven Rac^G12V^ expression caused a range of phenotypes, which we categorized as normal (Fig. 7B), intermediate (Fig. 7C), or severe (Fig. 7D). Normal egg chambers were indistinguishable from controls whereas the intermediate phenotype included nurse cell shrinkage, and the severe phenotype was characterized by nurse cell chromatin condensation and fragmentation (Fig. 7A-E), corresponding to early and late stages of apoptotic death observed in *slboGal4;UAS-Rac*^*G12V*^ egg chambers. The majority of *PG150-Gal4;UAS-Rac*^*G12V*^ egg chambers died prior to stage 10 (Fig. 7F). We conclude that Rac^G12V^ expression in stretch cells is sufficient to induce death through a similar series of steps as that observed in *slboGal4;UAS-Rac*^*G12V*^.

**Fig. 7.**
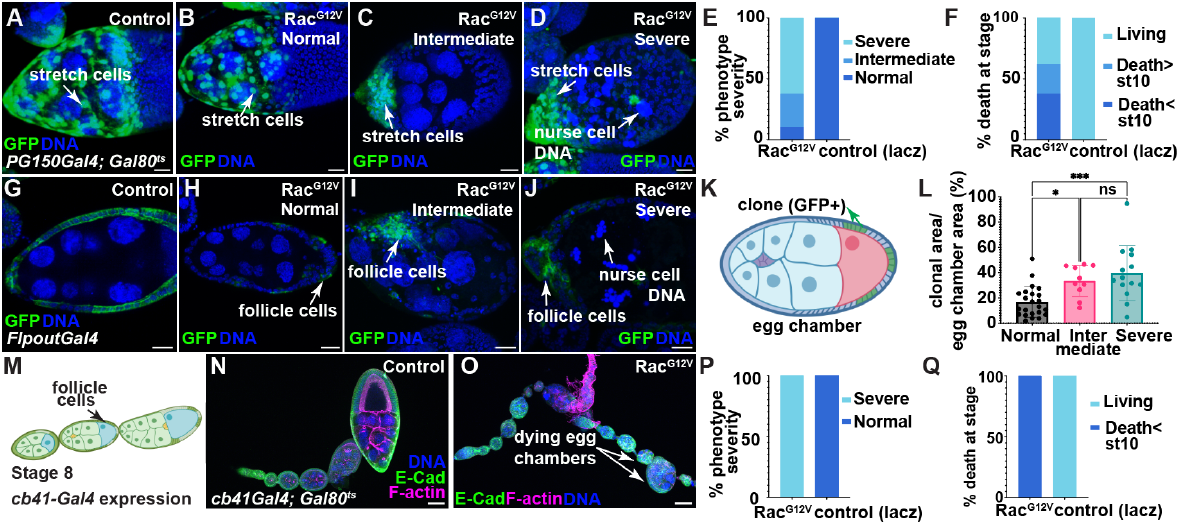
Large Rac^G12V^ follicle cell clones are required to kill the germline (A) Confocal image showing a control stage 10 egg chamber (*PG150;Gal80*^*TS*^,*UAS-GFP*.*nls; UAS-LacZ*). (B-C) Representative confocal images of egg chambers with *PG150;Gal80*^*TS*^, *UAS-GFP*.*nls,;UAS-Rac*^*G12V*^ depicting B. normal phenotype, C. intermediated phenotype, and D. severe phenotype of germline degradation. E-F. Plots of phenotype frequency (E) and germline death determined by chromatin condensation (F) (N=29). (G) Confocal image of a FlpOut Gal4 control egg chamber (H-J) Images of FlpOut Gal4 expression of Rac^G21V^ (GFP+ cells). (K) Schematic showing GFP clonal expression in the follicle cells. (L) Quantification of egg chamber death phenotype relative to clonal size (N=47). One way ANOVA with Tukey’s multiple comparison test was used to test statistical significance. ^*^ p<0.05, ^***^p<0.001. L. Schematic of *cb-41Gal4* expression. (M-N) Images of control or Rac^G12V^ expression with *cb-41Gal4*. (O-P) Plots of phenotype severity and stage of egg chamber death (N=22). Scale bars: 20 μm.

PG150 is expressed in ∼50 cells (37), much more than the 4-8 border cells that express slboGal4, so it was not clear if a similarly small subset of non-border cells expressing Rac^G12V^ would be capable of killing the germline. To investigate how many follicle cells expressing Rac^G12V^ were required to induce death, we used the FlpOut technique to drive Rac^G12V^ in clones of various sizes (Fig. 7G-J). We found that egg chamber death was more likely to occur when >30% of follicle cells expressed Rac^G12V^, as measured by the fraction of egg chamber area containing Rac^G12V^-expressing follicle cells (Fig. 7K-L). Rac^G12V^-expressing follicle cells also induced caspase activation in neighboring follicle cells (Fig. S5), demonstrating that Rac^G12V^-induced death does not require germline-specific sensitivity to apoptosis.

We also expressed Rac^G12V^ with *cb41-Gal4*, which is expressed in a mosaic pattern that includes many follicle cells [Fig. 7M, (38)]. Expression of Rac^G12V^ with *cb41-Gal4* induced death even in young egg chambers (Fig. 7N-Q). We conclude that Rac^G12V^-expressing follicle cells are sufficient to kill the germline but that a larger number of cells is required to match the potency of border cells. These results suggest that border cells are more efficient than other follicle cells at killing the germline.

## Discussion

Engulfment and killing of tumor cells by macrophages is a major mechanism by which CAR-macrophages and tumor-targeting antibodies such as rituximab and cetuximab exert antitumor activity (39–44). A significant challenge for these therapies is ensuring that macrophages achieve lethal clearance of sufficient tumor cells. FcγR^+^ myeloid cells instead perform sublethal trogocytic “nibbling,” which removes antigen– antibody complexes. This “antigen shaving” promotes immune escape rather than cancer cell death (45–51). Therefore elucidating the mechanisms that convert sublethal to lethal trogocytosis is of great interest. Sublethal nibbling and lethal whole cell engulfment are also frequent events in the normal development and homeostasis of multicellular organisms, including simple organisms like amoebae, worms, and flies as well as mammals including humans (50, 52). Therefore these are ancient cellular behaviors.

Rac is an ancient and deeply-conserved protein that serves as a critical node in the signaling and cytoskeletal pathways that govern chemotaxis and phagocytosis. Here we report cellular and molecular mechanisms by which Rac hyperactivation in a handful of border cells in the fruit fly ovary is sufficient to convert sublethal to lethal trogocytosis, thereby eliminating a much larger mass of cells. We delineate the steps by which border cells kill germline cells (Fig. 8): Rac hyperactivity drives biting off of fragments about twice as large as normal, which triggers apoptotic shrinkage in target cells followed by loss of the nuclear permeability barrier, caspase activation, and chromatin condensation and fragmentation. Ultimately all the follicle cells join in to consume the dying germline.

**Fig. 8.**
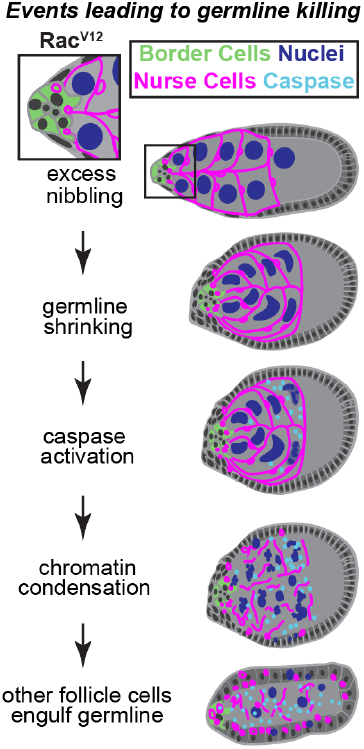
Schematic of steps in Rac^G12V^-mediated death. Border cells expressing Rac^G12V^ take bites out of nurse cells, which induces germline shrinkage, followed by caspase activation, loss of the nuclear permeability barrier, and chromatin condensation and fragmentation that spreads through the syncytium. As the germline dies, other follicle cells engulf and clear the dead germline cells.

We were surprised to find that border cells normally nibble on nurse cells as they migrate, in a Draper-and Rac-dependent manner, raising the question as to the purpose of this sublethal trogocytosis. Since Draper expression increases as border cells initiate both migration and nibbling, and Draper is required for border cells to complete their migration, nibbling may contribute to border cell motility or chemotaxis. Removal of chemoattractants from a path by migrating cells can produce self-generated gradients (53); however this is believed to occur through receptor-mediated endocytosis or enzymatic degradation rather than trogocytosis. Furthermore, the known chemoattractants for border cells are secreted rather than membrane-attached (54) (55), so nibbling would seem unnecessary to remove them. Another possibility is that nibbling clears space between nurse cells as it has been shown that even tiny spaces much smaller than a single border cell reduce the energy barrier for their movement (29). Alternatively, border cells may consume nurse cell material for fuel as has been described for cannibalistic cancer cells (6). Alternatively or in addition, trogocytosis may allow border cells to sense germline health. In zebrafish for example, macrophages nibble on hematopoietic stem and progenitor cells to assess their health, and if the phagocytes sense oxidative stress they phagocytose and remove the stressed progenitor (56). In *C. elegans* development, endodermal cells engulf large, mitochondria-rich lobes from primordial germ cells, presumably to prune unfit organelles (57). Similar cell-cell interactions may be more widespread than previously appreciated, serving context-dependent functions. As in border cell nibbling, Rac1 is a central molecular player in sublethal and lethal trogocytosis as well as whole cell engulfment in most if not all of these examples (52).

The work presented here shows that there are multiple, related yet distinct mechanisms by which follicle cells can engulf and/or kill the germline. We find that Rac-mediated germline killing shares similarities to both starvation mediated germline engulfment and developmental engulfment of nurse cells and yet is distinct from both. Rac-mediated death is initiated by border cell trogocytosis of nurse cells, whereas starvation-induced death starts with germline apoptosis, and developmental death begins when stretch follicle cells enwrap and acidify individual nurse cells.

Our data indicate that constitutively activated Rac engages multiple pathways to promote germline killing. Rac is required for the formation of F-actin–rich protrusions that generate the phagocytic cup, and Rac hyperactivity enables border cells to take larger bites.

We previously reported that *draper* mutants partially rescue Rac-mediated germline death: whereas ∼70% of *slboGal4;UAS-Rac*^*G12V*^ egg chambers die in the presence of Draper, only 35% die in homozygous *drpr* mutants (15). The rescue appeared to prevent even the initiation of death, so it is likely that Draper expression in border cells enhances their ability to take the larger bites required to convert sublethal to lethal trogocytosis. Prior work shows that Draper promotes germline death by inducing oxidative stress (34) and driving acidification-mediated nurse-cell degradation during development (26). Rac is also known to activate NADPH oxidase to generate the oxidative burst in phagocytes, so Rac and Draper may synergize to generate oxidative stress in addition to physical damage. We only observed germline acidification late in the death process, after upregulation of Draper expression in all follicle cells. Thus, only after nurse cells die, do Rac^G12V^-expressing egg chambers engage processes similar to those that are normally employed during stage 12.

Rac-induced death resembles starvation-induced death in that it depends on caspase-mediated apoptosis, the oocyte is not spared, and all follicle cells eventually engulf the dying germline. These similarities might arise from the egg chamber stage since Rac1^G12V^-expressing border cells kill the germline in stages 8-9, which is similar to starvation-induced death at stages 7-8. However unlike starvation-mediated death but like developmental death, follicle cells initiate Rac^G12V^-induced killing non-autonomously in a process that requires engulfment genes. In starvation-induced death, caspase-dependent germline apoptosis precedes engulfment. Intriguingly, treatment of *Drosophila* with rapamycin, which inhibits the nutrient sensor Target of Rapamycin (TOR), induces follicle cells to engulf the germline (58)), and this effect is conserved in mammalian oogenesis. Thus, follicle cells can sense nutrient stress, either autonomously or nonautonomously, and may be primed to kill and engulf the germline under suboptimal conditions. However, the observation that Rac^G12V^ induces caspase activation in neighboring follicle cells indicates that Rac^G12V^-induced death does not require a germline-specific sensitivity to apoptosis. It is interesting that Rac^G12V^-expressing border cells engulf but do not kill polar cells (15), even when they are stretched and split, including obvious nuclear deformation, suggesting that polar cells are especially resistant to apoptosis. Identifying the tipping points, such as levels of Rac activity in the engulfing cell and vulnerabilities in the targets, that make the difference between sublethal and lethal phagocytic behavior enhance both our understanding of homeostatic checkpoints and the ability to engineer therapeutic phagocytes to remove unwanted cells.

## Materials and Methods

### Fly husbandry and genetics

Flies were maintained in cornmeal-yeast food. Crosses were kept at 18 °C for stocks with *Gal80*^*TS*^ while other crosses were kept at room temperature or 25 °C and transferred to fresh food every 2-3 days. Control and experimental conditions were kept at the same temperature for each experiment. Prior to ovary dissections, 5-7 female flies and 1-2 males were incubated together with supplemental yeast at 29 °C overnight.

### Live imaging

Whole ovaries were first dissected with forceps from flies in live imaging media [Schneider’s media (Thermo Fisher Scientific) supplemented with 20% heat-inactivated Fetal Bovine Serum (Sigma-Aldrich) and 1x antimycotic/antibiotic (VWR) at pH 6.85-6.95] in a depression slide as described (59). Then, individual ovarioles were teased from the muscle sheath by holding the germarium using forceps. Dissected ovariole strands were resuspended in live imaging media with 0.4 mg/mL bovine insulin (I1882, Sigma-Aldrich) and 0.1 % agarose. For lysotracker and dextran staining, ovaries were incubated at 1:1000 immediately prior to live imaging. For caspase inhibition, ovarioles were incubated in 100 μM Q-VD-OPh (APExBio) in live imaging media for 30 minutes on a rocker prior to imaging. BAPTA-AM, cell permeant Ca2+ chelator (ThermoFisher) was incubated for 45 minutes on a rocker prior to live imaging at a concentration of 50 micromolar 0.02% with Pluronic F-127 (Invitrogen). MitoSOX Red (Thermofisher) was added at a concentration of 5uM and incubated for 10 mins with resuspension every 5 minutes. Xrhod-1(Invitrogen) was added at a concentration of 5 micromolar. These dyes and inhibitors were maintained in the culture media throughout imaging. A 20×20 mm coverslip (No. 1.5 Fisherbrand) was split in half to provide space between the imaging coverslip and the sample, and 4 uL of live imaging media was used to stabilize each part of the coverslip 100 microliters of media with the egg chambers was loaded into the center of the split 20x 20 cover slips on a Lumox dish (Sarstedt). A larger cover slip (40X20 mm No. 1.5 Fisherbrand) was then placed over the sample. The sample was inverted to allow samples to settle close to the coverslip. After 5 minutes, 20 uL of halocarbon oil was added around the periphery of the coverslip.

Samples were imaged on a ZEISS LSM 800 or ZEISS 980. Images shown in Figure 3A-E were acquired on a ZEISS LSM 780 in Lambda mode and images were processed by spectral unmixing to distinguish fluorescent markers with overlapping emissions spectra.

### Immunostaining

Ovarioles were dissected from ovaries in live imaging media, allowed to settle, and fixed in 4% paraformaldehyde in 1X PBS for 15 minutes at room temperature. Ovarioles were washed 3 times with 1x PBST at room temperature, and then incubated in primary antibodies overnight (Draper (DSHB, 1:50), DCAD2, (DSHB, 1:15), pJNK (Cell Signaling Technologies 1:200), cDcp-1 (Cell Signaling Technologies, 1:200) at 4°C. The next day, ovarioles were washed thrice with 1 X PBST and incubated with secondary antibodies at 1:200 (Goat anti-Rat IgG [H+L] Cross-Adsorbed Secondary Antibody, Alexa Fluor 647, 488, or 568, Invitrogen, Goat anti-Mouse IgG [H+L] Cross-Adsorbed Secondary Antibody, Alexa Fluor 488 and 568; Invitrogen) at room temperature. Samples were also incubated with phalloidin-Atto-647N (1:500; Sigma-Aldrich), and Hoechst (1:1000 of 10 mg/mL stock, Invitrogen). Samples were incubated with secondary antibodies, phalloidin and Hoechst together for 1 hour at room temperature. Ovaries were washed with 1X PBST three times then suspended in Vectashield (H-1000; Vector Laboratories) and incubated overnight at 4°C before mounting onto slides.

### Image analysis

#### Egg chamber death scoring

In fixed imaging, death was scored per ovariole, i.e. if there was one or more dead egg chamber(s) at the end of an ovariole strand. An egg chamber was scored as dead when the germline chromatin (DNA labeled with Hoescht) became condensed and/or fragmented. In live imaging of stage 9 egg chambers, death was scored when the germline chromatin, labeled with histone RFP, underwent condensation and/or fragmentation prior to the end of the imaging session (within 4.5 hours).

#### Nurse cell membrane analysis

Nurse cell membrane area was analyzed from a single z-stack or live egg chambers. The area of nurse cell membrane inside the border cells was measured by drawing an ROI around the nurse cell membrane fragment in the border cells, using *slbo*4X-PH-EGFP or follicle cells in Image J. The largest membrane fragment is shown for each egg chamber.

#### Germline area measurements

The entire area of the nurse cells were traced in Image J using the DIC images from movies with UbiHis-RFP and *slbo*4XPH-EGFP every 25 minutes until the germline died (indicated by chromatin condensation) or moved excessively or of the field of view. The average change in area per 25 minutes is reported for each egg chamber.

#### Migration Index

To measure migration indices, the location of the border cell cluster was assessed in stage 10 egg chambers, defined as when the oocyte occupies ½ of the egg chamber volume and the anterior border of the columnar follicle cells aligns with the anterior oocyte border. The position of the border cell cluster was manually scored in the following categories: 0%-still attached at the anterior end of the egg chamber; <50% - detached but located in the anterior half of the migration path; >50% - within the posterior half of the migration path; and 100% - at the oocyte border.

#### Fluorescence reporters and intensity assays

To measure the fluorescence intensity of Draper border cells at different stages, a small box (10 x 10 pixels) was drawn around the border cell cluster, and an equivalent box was selected outside of the egg chamber for background subtraction, for each image. For pJNK, a 20 x 20 pixel box was drawn around a GFP+ cell and a GFP-cell within the same egg chamber, and an equivalent box was selected outside of the egg chamber for background subtraction. The ratio was calculated by the (mean fluorescence intensity GFP+ cell - background)/(mean fluorescence intensity of the GFP-cell).

The percentage of cDcp-1 positive and *puc*-LacZ positive cells scoring were scored manually. cDcp-1 was considered positive if it accumulated within the germline (usually observed at the periphery of the nurse cells, which is consistent with prior reports using this staining. *puc*-LacZ was considered positive if there was a detectable enrichment of the signal within the nucleus.

#### Data representation and statistical analysis

All sample sizes and middle bars are reported in the figure legends. For analysis the inclusion criteria were all undamaged samples, correct egg chamber stage, and in-focus images. Exclusion criteria include damaged, out of focus, or incorrect stage time points and images. For live imaging, all stage 9 egg chambers were included unless they went out of view or focus due to movement during the imaging or if they became abnormally flattened during the imaging session. Statistical tests used are reported in the figure legends ^*^ indicates p<0.05, ^**^ p<0.01. ^***^ p<0.001, ^****^ p<0.0001. Image processing differences are noted in figures and legends but were generally processed the same way as controls.

## End Matter

## Acknowledgments

We would like to thank the Bloomington Drosophila Stock Center for fly stocks used in this paper and Flybase for its invaluable service and resources. We would like to thank Dhruv Kaushik, Emory Campbell and Wesley Kim for their technical assistance on this project. We would like to thank the members of the Morrissey and Montell labs for their feedback on this work. This work was supported by the NIH grant R01GM06425 to D.J.M. L.P. was funded by the American Cancer Society (PF-22-091-01-MM).

## Author Contributions

A.K.M. designed research, performed research, and analyzed data. L.P. designed research, performed research, analyzed data, and wrote the paper. M.S. performed research and analyzed data. D.J.M. designed research, analyzed data, and wrote the paper. All authors have carefully reviewed and approved the final version of the manuscript.

---

**Fig. S1.**
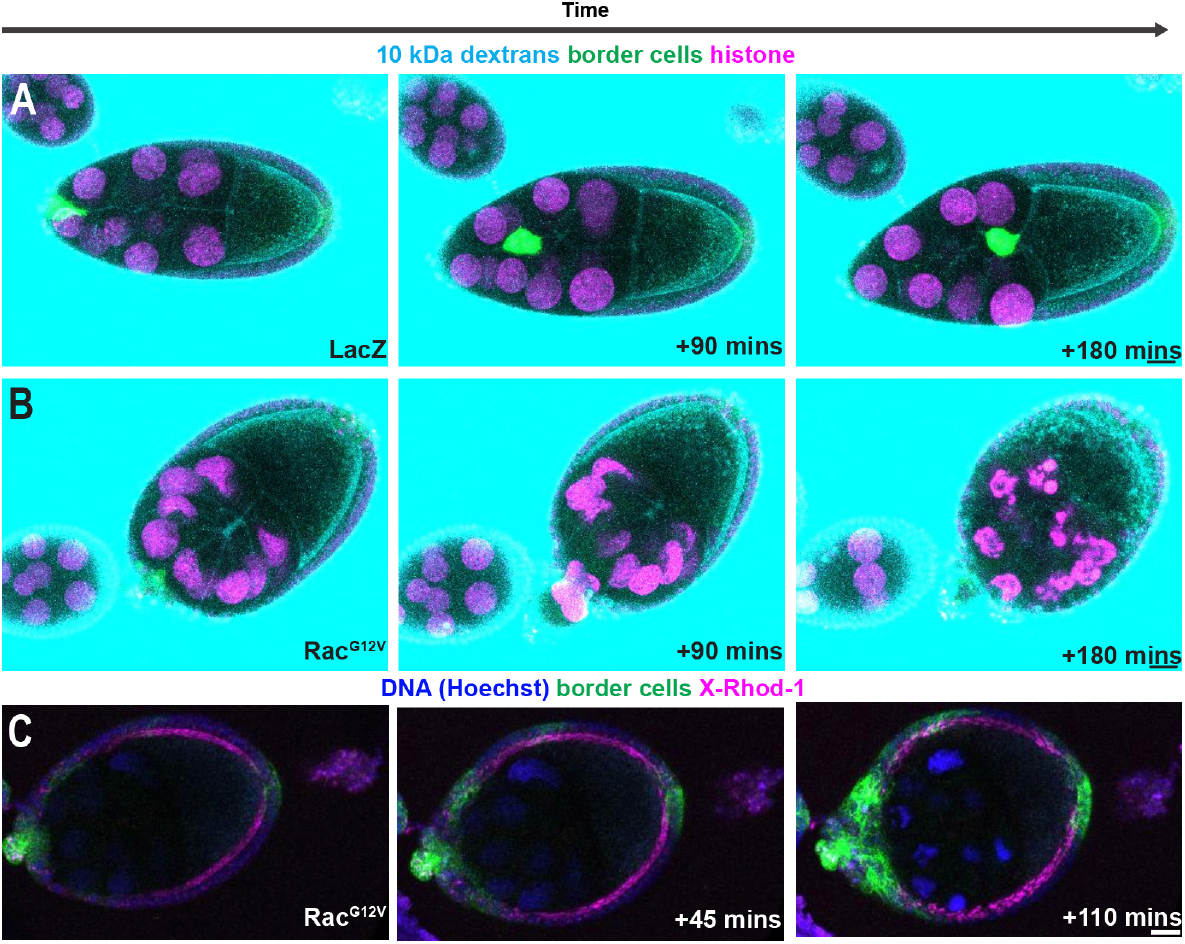
Germline cells do not have detectable breaches in the plasma membrane prior to chromatin condensation. (A-B) Images from a timelapse series of egg chambers overexpressing *UAS-LacZ* control(A) or *UAS-Rac*^*G12V*^(B) with *slboGal4*. Border cells are marked with *slbo*4X-PH-EGFP (green). Nuclei are labeled with UbiHis-RFP (magenta). Egg chambers are incubated with 10 kDa dextrans conjugated with Alexa 647 (cyan). (C) Images from a timelapse series of an egg chamber expressing *slboGal4;UAS-Rac*^*G12V*^ *;UAS-PH(PLCδ)-EGFP* (green) and stained with XRhod-1, a calcium indicator (magenta), and Hoechst to label DNA (blue). Scale bars: 20 μm.

**Fig. S2.**
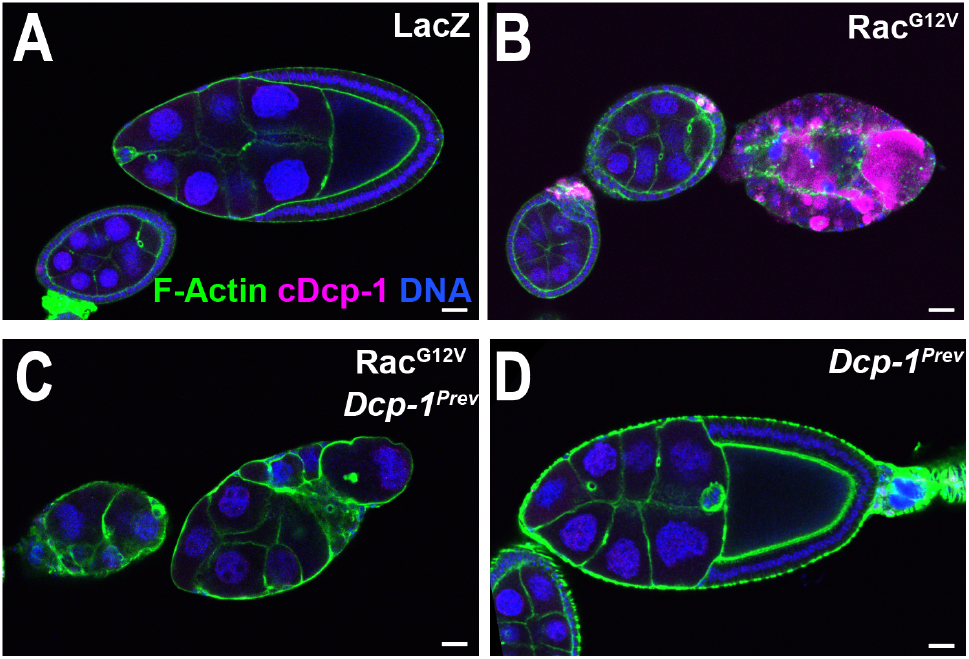
Suppression of nurse cell chromatin condensation by *Dcp-1*^*Prev*^. (A-D) Confocal images of egg chambers from *c306Gal4; Gal80*^*TS*^ flies stained for cleaved caspase (cDcp-1, magenta), F-actin (green), and DNA (blue) expressing *UAS-LacZ* (A), *UAS-Rac*^*G12V*^ (B), or *UAS-Rac*^*G12V*^ in a homozygous *Dcp-1*^*Prev*^ mutant (C). D shows a homozygous *Dcp-1*^*Prev*^ mutant egg chamber without hyperactive Rac.

**Fig. S3.**
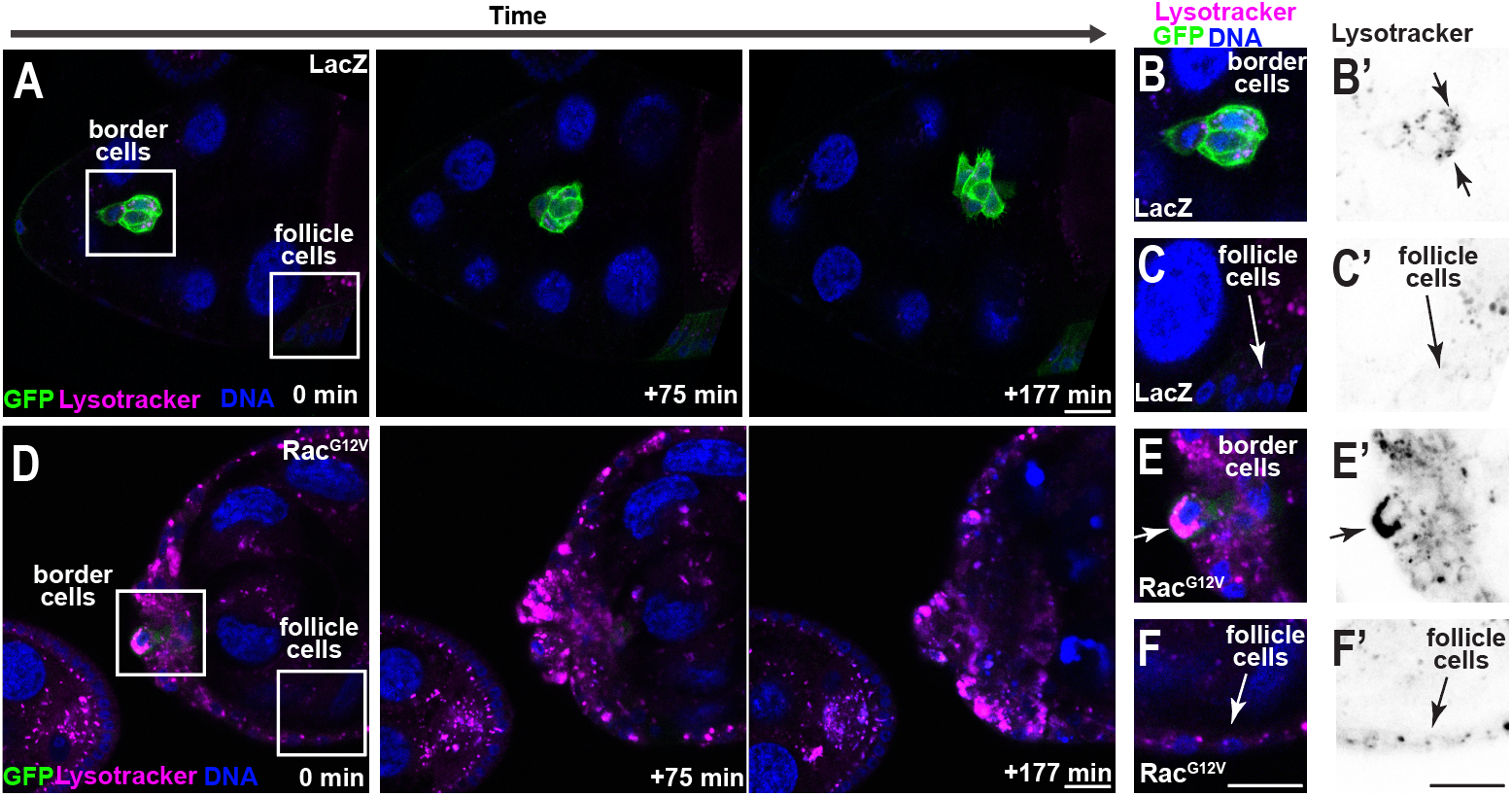
Lysotracker accumulates in Rac^G12V^ border cells and other follicle cells prior to germline death. (A-C) Images from a timelapse series of *slboGal4;UAS-LacZ A*) expressing UbHisRFP (blue), *slbo*4x-PH-EGFP (green) and stained for lysotracker-deep red (magenta). (B-B’) Magnified image at time=0 minutes highlights lysotracker puncta in the merged image of border cells (B)’ and inverted fluorescence of lysotracker (B’). (C) Magnified image of follicle cells in *slboGal4;UAS-LacZ* at time=0 mins. (C’) Inverted fluorescence of lysotracker. Arrows indicate follicle cells. (D-F) Images from a timelapse series of a *slboGal4;UAS-Rac*^*G12V*^ (D). (E) Magnified image of Rac^G12V^ border cells at 0 minutes. (E’) inverted fluorescence of lysotracker. Arrow indicates accumulation of lysotracker. (F) Magnified image of follicle cells in *slboGal4;UAS-Rac*^*G12V*^ egg chambers at time=0 mins. (F’) inverted fluorescence of lysotracker. Arrows indicate follicle cells. See also Movies S14-15.

**Fig. S4.**
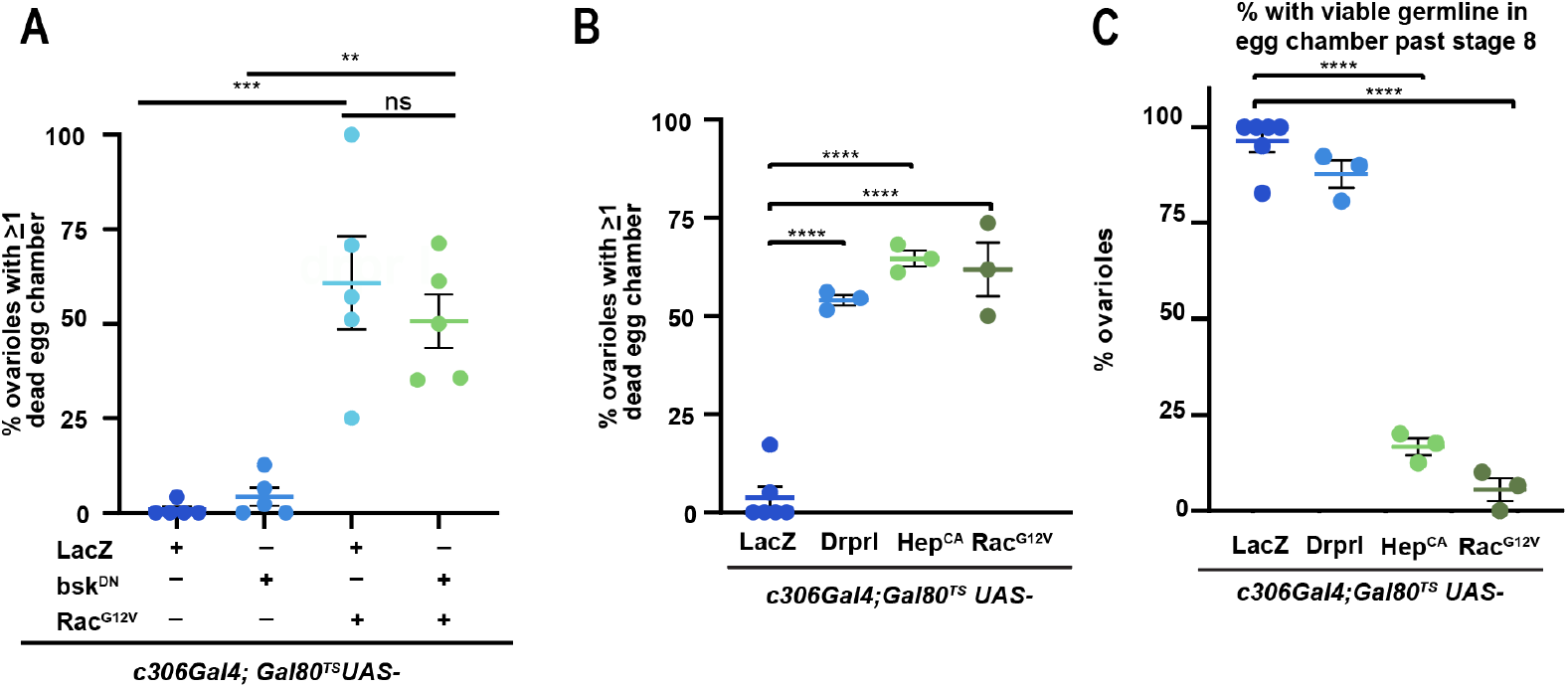
Expression of Rac^G12V^, Draper, or hepCA with border cell driver, *c306Gal4; Gal80*^*TS*^ induces egg chamber death. (A-B) Plot showing mean +/-SEM frequencies of percent of ovariole strands with germline death for the indicated conditions. (C) Plot showing the mean +/-SEM frequencies of ovariole strands with viable egg chambers past stage 8. Each dot represents an experimental replicate analyzing n>18 egg chambers P values are from a One Way ANOVA with Post-Hoc Tukey. ^****^p<0.0001, ^**^p<0.01, ^***^p<0.001, ns: not significant

**Fig. S5.**
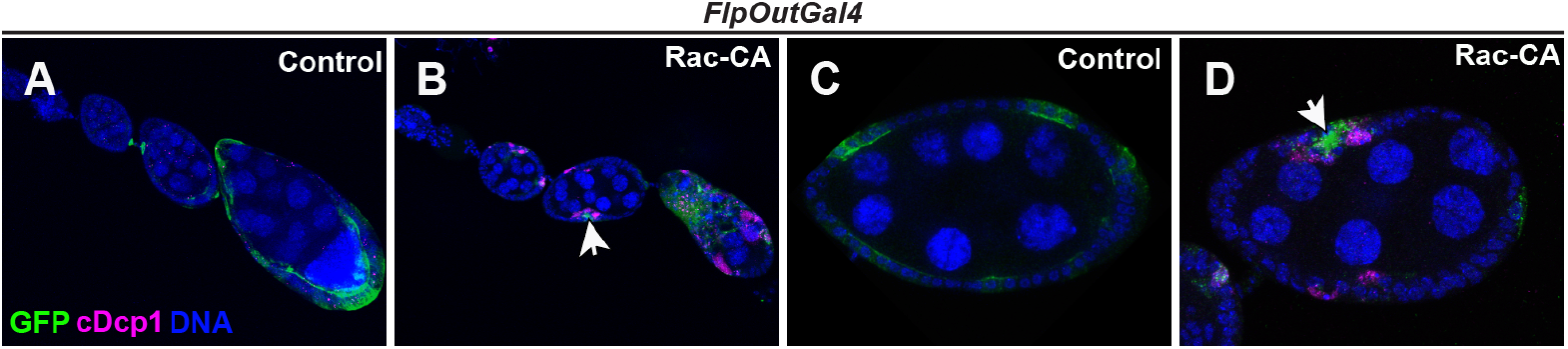
Rac^G12V^ follicle cell clones kill neighboring cells. (A-D) Confocal images of ovarioles expressing *hsFlpAyGal4 UAS-GFP* along with *UAS-LacZ* (A,C) or *UAS-Rac*^*G12V*^ (B,D) stained with Hoechst to mark DNA (blue) and cleaved Dcp-1 (cDcp-1) to mark active caspase (magenta). Arrows indicate follicle cell death.

**Supplementary Table 1.**
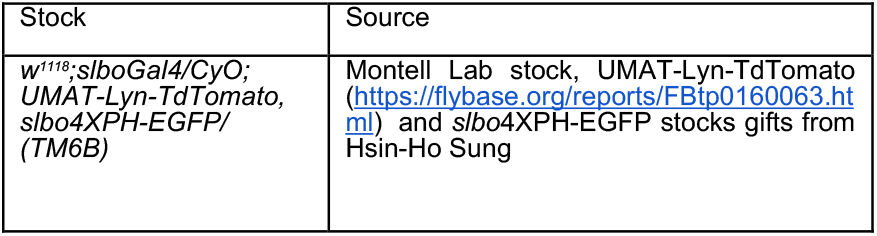
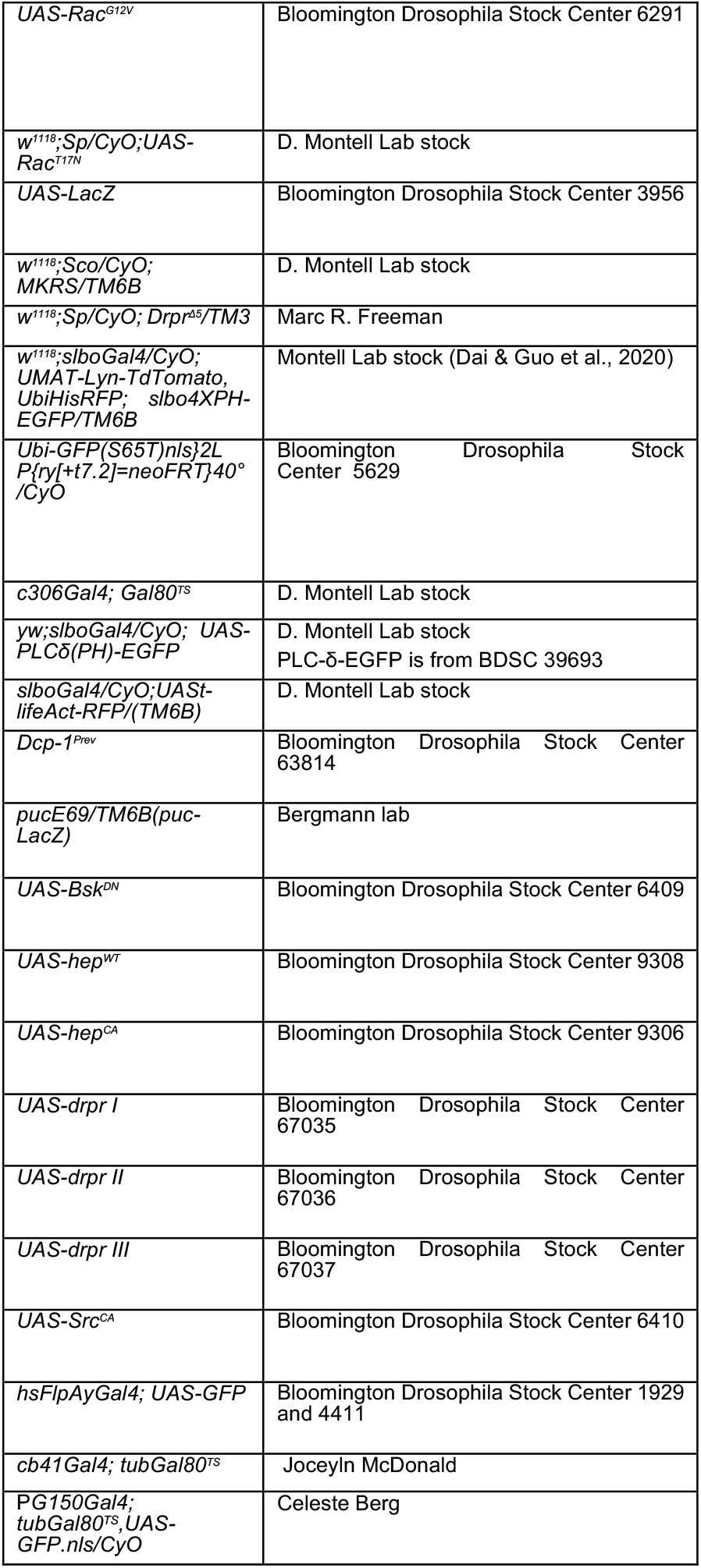
Drosophila Fly Stocks.

## Supplemental Movies

Movie S1.

Control border cells accumulate puncta of nurse cell membranes. *slboGal4; UAS-LacZ*; *slbo*4XPHEGFP(green, border cell membranes) and UMAT-LynTdTomato (nurse cell membranes, magenta). Images were acquired every 3 minutes, single z-slices are shown. Scale bar: 20 μm.

Movie S2.

Rac^G12V^-expressing border cells take in larger pieces of nurse cell membranes. *slboGal4; UAS-Rac*^*G12V*^; *slbo4XPHEGFP* and UMAT-LynTdTomato. Images were acquired every 3 minutes, single z-slices shown. Scale bar: 20 μm.

Movie S3.

Control border cell migration. Egg chambers expressing *slboGal4; UAS-LacZ, slbo4*XPH-EGFP (green), UMAT-LynTd-Tomato (Cyan), and Ubi-Histone-RFP (magenta). Images were acquired every 3 minutes, max z projections are shown. Scale bar: 10 μm.

Movie S4.

Germline death induced by expression of *slboGal4; UAS-Rac*^*G12V*^. Egg chambers also expressed *slbo*4XPH-EGFP (green), UMAT-LynTd-Tomato (Cyan), and Ubi-Histone-RFP (magenta). Images were acquired every 3 minutes, max z projections are shown. Scale bar: 10 μm.

Movie S5.

Higher magnification example of control border cell migration. Egg chambers expressing, *slboGal4; UAS-LacZ, slbo*4XPH-EGFP (green), UMAT-LynTd-Tomato (Cyan), and Ubi-Histone-RFP. Images were acquired every 3 minutes, max z projections are shown. Scale bar: 20 μm.

Movie S6.

Higher magnification example of *slboGal4; UAS-Rac*^*G12V*,^. *slbo*4XPH-EGFP (green), UMAT-LynTd-Tomato (Cyan), and Ubi-Histone-RFP (magena), Images were acquired every 3 minutes,max z projections are shown. Scale bar: 20 μm.

Movie S7.

Second higher magnification example of *slboGal4; UAS-Rac*^*G12V*,^ *slbo*4XPH-EGFP. (green), UMAT-LynTd-Tomato (Cyan), and Ubi-Histone-RFP (magena), Images were acquired every 3 minutes and max z projections are shown. Scale bar: 20 μm.

Movie S8.

Control egg chambers retain the nuclear permeability barrier during stage 9. Egg chamber expressing Ubi-GFP.NLS (inverted grayscale fluorescence) Images were acquired every 5 minutes, and MAX z projection is shown. Scale bar: 20 μm.

Movie S9.

Posterior germline cell nuclei lose the permeability barrier, and the loss spreads through the syncytium during death. Egg chamber expressing slboGal4; *UAS-Rac*^*G12V*^ *slbo*4XPH-EGFP Ubi-GFP.NLS (inverted grayscale fluorescence) Images were acquired every 5 minutes, and MAX z projection is shown. Scale bar: 20 μm.

Movie S10.

Egg chambers grow during stage 9. Egg chambers expressing, *slboGal4; UAS-LacZ, slbo*4XPH-EGFP (green), UbiHisRFP (magenta). The differential interference contrast (DIC) image is overlaid with fluorescent channels. Images were acquired every 5. Minutes, and single z-slices are shown. Scale bar: 20 μm.

Movie S11.

Egg chambers shrink prior to Rac^G12V^ mediated chromatin condensation. Time-laspe series from egg chambers expressing slboGal4; *UAS-Rac*^*G12V*^ *slbo*4XPH-EGFP (green), UbiHisRFP (magenta). The differential interference contrast (DIC) image is overlaid with fluorescent channels. Images were acquired every 5 minutes, and single z-slices are shown. Scale bar: 20 μm.

Movie S12.

QVD rescues Rac^G12V^ mediated chromatin condensation. Timelaspe series from egg chambers expressing *slboGal4; UAS-Rac*^*G12V*^ *slbo*4XPH-EGFP (green), UbiHisRFP (magenta) and treated with 100 micromolar QVD for 30 minutes prior to imaging. The differential interference contrast (DIC) image is overlaid with fluorescent channels. Images were acquired every 5 minutes, and single z-slices are shown. Scale bar: 20 μm.

Movie S13

BAPTA-AM treatment of *slboGal4; UAS-Rac*^*G12V*^ egg chambers. Timelaspe series from egg chambers expressing *slboGal4; UAS-Rac*^*G12V*^ *slbo*4XPH-EGFP (green), UbiHisRFP (magenta) treated with 50 micromolar BAPTA-AM. The differential interference contrast (DIC) image is overlaid with fluorescent channels. Images were acquired every 5 minutes, and single z-slices are shown. Scale bar: 20 μm.

Movie S14.

Control border cells have small lysotracker puncta during migration. Timelapse series of a (*slboGal4;UAS-LacZ*) expressing UbHisRFP (blue), *slbo*4PHEGFP (green) and stained for lysotracker-deep red (magenta). Images were acquired every 3 minutes, and a max z-projection is shown. Scale bar: 20 μm.

Movie S15.

Border cells and follicle cells accumulate lysotracker during Rac-mediated death. Timelapse series egg chambers expressing *slboGal4;UAS-Rac*^*G12V*^, UbHisRFP (blue), and *slbo*4PHEGFP (green) that are stained for lysotracker-deep red (magenta). Images were acquired every 3 minutes, and a max z-projection is shown. Scale bar: 20 μm.

Movie S16.

MitoSox staining (magenta) in control egg chambers expressing slboGal4; *UAS-LacZ;UAS-PH(PLCδ)-GFP* (green). Images were acquired every 3 minutes.

Movie S17.

MitoSox staining (magenta) in egg chambers expressing *slboGal4;UAS-;Rac*^*G12V*^*;UAS-PH(PLCδ)-GFP* (green). Images were acquired every 3 minutes.

